# gcFront: a tool for determining a Pareto front of growth-coupled cell factory designs

**DOI:** 10.1101/2021.10.12.464108

**Authors:** Laurence Legon, Christophe Corre, Declan G. Bates, Ahmad A. Mannan

**Affiliations:** Warwick Integrative Synthetic Biology Centre, School of Engineering, University of Warwick, Coventry, UK; Warwick Integrative Synthetic Biology Centre, School of Life Sciences, University of Warwick, Coventry, UK

## Abstract

**Motivation:** A widely applicable strategy to create cell factories is to knock out (KO) genes or reactions to redirect cell metabolism so that chemical synthesis is made obligatory when the cell grows at its maximum rate. Synthesis is thus growth-coupled, and the stronger the coupling the more deleterious any impediments in synthesis are to cell growth, making high producer phenotypes evolutionarily robust. Additionally, we desire that these strains grow and synthesise at high rates. Genome-scale metabolic models can be used to explore and identify KOs that growth-couple synthesis, but these are rare in an immense design space, making the search difficult and slow.

**Results:** To address this multi-objective optimization problem, we developed a software tool named gcFront - using a genetic algorithm it explores KOs that maximise cell growth, product synthesis, and coupling strength. Moreover, our measure of coupling strength facilitates the search so that gcFront not only finds a growth coupled design in minutes but also outputs many alternative Pareto optimal designs from a single run - granting users flexibility in selecting designs to take to the lab.

**Availability and Implementation:** gcFront, with documentation and a workable tutorial, is freely available at GitHub: https://github.com/lLegon/gcFront, the repository of which is archived at Zenodo, DOI: 10.5281/zenodo.6338595 (Legon *et al*., 2022).

**Supplementary Information:** Supplementary notes and data files are available at Bioinformatics online.

## INTRODUCTION

Genome-scale constraint-based models (GSMs) are used to explore gene or reaction knockouts (KOs) that redirect cell metabolism to chemical overproduction (Maia *et al*., 2016). A promising strategy for enabling robust production seeks KO combinations that couple chemical synthesis with cell growth so that it is made obligatory at maximum growth rate (Feist *et al*., 2010). KOs can disrupt metabolism to result in poorer performance than predicted, but growth coupling enables the selection of higher producing phenotypes by selecting faster growing cells through adaptive laboratory evolution (ALE). KOs by gene deletion are easily implemented in the lab, and since they remain fixed in the face of evolution, as opposed to engineering changes in gene expression, ALE has been shown to find strains with synthesis and growth rates near the optimal values predicted from GSMs (Tokuyama *et al*., 2018). However, if the coupling is weak, cells will not synthesise the product unless they grow close to their theoretical maximum. Instead, KOs that create a strong coupling result in evolutionarily robust phenotypes with robust synthesis, and so are particularly appealing. Specifically, stronger coupling will strongly impair growth for small impediments in product synthesis, so higher producers will be reselected over evolutionary time, and it also helps conserve synthesis rates even if cells grow at suboptimal rates, for instance in large fermenters (Supplementary Fig. 1). In addition to strong coupling, we also desire that these strains grow fast but also synthesise rapidly. Identifying the KO sets, i.e. designs, that maximise these criteria is a multi-objective optimization problem. However, there are inherent trade-offs between some of these objectives, so solving this problem will give a set of alternative optimal designs where for each design each objective cannot be improved without sacrificing some of the others. This is known as a Pareto front of optimal designs. Multi-objective optimization has been applied in metabolic engineering, for instance to kinetic models to find Pareto optimal reaction kinetics that maximize synthesis (Sendín *et al*., 2006; Vera *et al*., 2003), and tools have been developed for use on GSMs to determine genetic manipulations to maximize growth and synthesis (Andrade *et al*., 2020; Patané *et al*., 2019). Other tools have been developed to find growth-coupled designs (Feist *et al*., 2010; Ohno *et al*., 2014; Alter and Ebert, 2019), yet there is no tool to determine optimal designs that maximise coupling strength, growth, and synthesis, in order to create evolutionarily robust strains with high productivity and robust synthesis – critical for industrial application. Moreover, though growth coupling is a widely applicable strategy (von Kamp and Klamt, 2017) KOs enabling this are rare, making the search for them difficult and slow (Ohno *et al*., 2014). To address this key gap and problem, we developed a user-friendly software tool named gcFront that uses a genetic algorithm to search for KOs that maximize these three objectives, for any chemical and host of interest. Moreover, our proposed measure of coupling strength facilitates the search through the design space, so a run of gcFront outputs many Pareto optimal designs in reasonable timeframes.

## THE gcFRONT WORKFLOW

gcFront works in MATLAB, with dependencies on the COBRA toolbox (Heirendt *et al*., 2019) for analysis of a compatible GSM; and the MATLAB Global Optimization toolbox for solving the multiobjective optimization problem (Supplementary Note 1A). The workflow, detailed in Supplementary Note 1B and Supplementary Fig. 2, entails four fundamental steps.

### Inputs

Two interactive windows allow the user to define the GSM and target metabolite product or its exchange reaction, and optional inputs (Supplementary Table 1), such as maximum number of KOs and search time.

### Pre-processing

To reduce the search space of reactions, gcFront automatically identifies and removes dead reactions, lumps unbranched pathways into composite reactions, and excludes *in silico* essential single KOs for growth or synthesis.

### Solving the optimization problem

To determine growth-coupled designs, gcFront solves the multiobjective optimisation problem defined in Supplementary Note 2A. Our measure of coupling strength shapes the search landscape; it defines weak and strong coupling but also distinguishes between uncoupled designs (Supplementary Note 2B, Supplementary Fig. 3a). It assigns higher values to KOs that reduce the cost to growth for increases in the maximum allowable synthesis, thus driving a bias to gc-designs (Supplementary Fig. 3b and 3c) to ease the search.

### Post-processing and output

On termination (conditions in Supplementary Table 1), many Pareto optimal KO sets are found from a single run. Some proposed designs may contain redundant KOs, so to minimise the number of KOs of each design any KO that can be removed from those designs without any loss in performance is removed. The Pareto front of all designs (KOs) and their performance is then output to an interactive plot, a table in the command window, and a .csv file. Users can select designs they deem suitable for their chemical and host of interest, based on bespoke combinations of the performance metrics. A tutorial is given in Supplementary Note 3.

## COMPARATIVE PERFORMANCE ASSESSMENT

To test gcFront’s performance, we compared it to other MATLAB-based procedures that identify growth-coupled (gc-)designs, including RobustKnock (Tepper and Shlomi, 2009) as implemented in OptPipe (Hartmann *et al*., 2017); gcOpt (Alter and Ebert, 2019); FastPros (Ohno *et al*., 2014); and OptGene (Patil *et al*., 2005) as implemented in COBRA (Heirendt *et al*., 2019). We ran each for 6 hours and saved the gc-designs found and the time they needed to find their first gc-design, while repeating this three times for gcFront and OptGene because of the stochastic nature of searching with a genetic algorithm. For a fair comparison, we ran each algorithm using the *E. coli* GSM model iML1515 (Monk *et al*., 2017), for non-essential reaction KOs (*in silico* and based on (Goodall *et al*., 2018)), for synthesis of succinate, tyrosine, and pyruvate, as example products (detailed in Supplementary Notes 4 and Fig. 1). gcFront found the first gc-design in 38% less time than gcOpt for succinate synthesis, 98% less time than RobustKnock for tyrosine synthesis, and orders of magnitude less time than the other methods and products (Fig. 1a, Supplementary Data). Its power was especially apparent when searching for designs of tyrosine and pyruvate synthesis – still finding designs in minutes despite these designs, of at least six KOs, being rarer vs three KOs found for succinate (Supplementary Fig. 4). Furthermore, though the single gc-design found with other methods lay near the Pareto front of gc-designs from gcFront, gcFront offered many designs that achieved at least higher coupling strength (Fig. 1b, Supplementary Data).

**Fig. 1.**
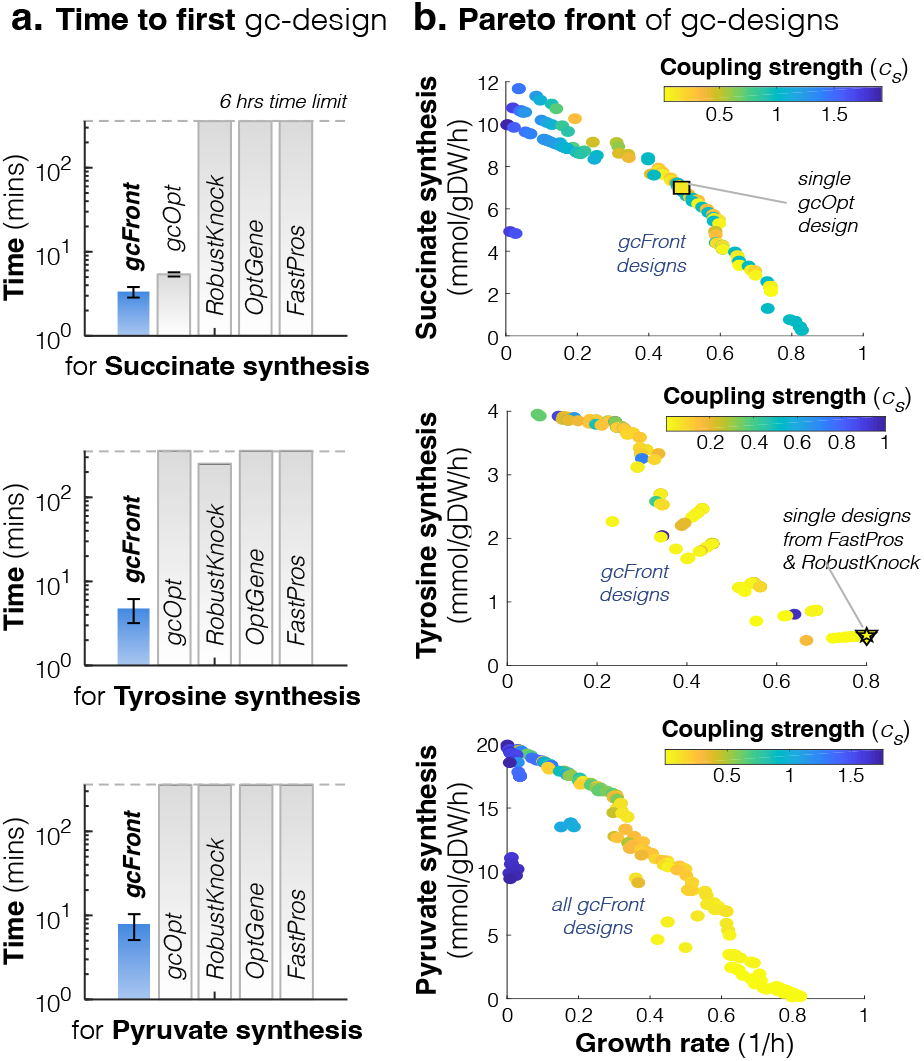
gcFront finds many Pareto optimal growth-coupled designs, faster, and with superior performance vs other algorithms. The speed and designs found from 6-hour runs of gcFront were compared to those of RobustKnock, gcOpt, FastPros, and OptGene (see Supplementary Note 4) on a MacBook Pro (2.3 GHz Quad-Core Intel core i5 processor, 8GB 2133 MHz LPDDR3 RAM). Designs were based on KOs of only non-essential, gene-associated reactions, for the synthesis of three example products: succinate, tyrosine, and pyruvate from the *E. coli* iML1515 GSM model, in aerobic, minimal media with glucose. (**a**). Time to identify the first gc-design from each procedure. Due to the stochastic nature of searching using the genetic algorithm in OptGene and gcFront, the average (bars) and standard deviation (error bars) of times are reported from three runs (N=3, ±SD). (**b**). Pareto fronts of all gc-designs found from three 6-hour runs.

## DISCUSSION

gcFront can find a multitude of Pareto optimal growth-coupled designs for evolutionarily robust cell factories, from a single run, in a computationally efficient manner. With the key input being the genome-scale metabolic network model of the cell host with the biochemistry of the engineered product synthesis pathway, gcFront should be widely applicable for designing growth-coupled synthesis of any compound, from any host, driving the design step in the design-build-test-learn cycle (Carbonell *et al*., 2018). Since each design provides a different balance between the maximised objectives, the user has the flexibility to select designs with the balance they deem most suitable to the cell host and chemical product of interest, e.g. sacrifice growth for stronger coupling and synthesis, for instance for more robust pyruvate synthesis (Fig. 1b); or sacrifice synthesis for higher growth and stronger coupling, for instance for higher volumetric productivity with robust synthesis of succinate (Fig. 1b) – making it widely applicable to different contexts. gcFront is also user friendly, but versatile – the interactive user interface means no coding is required, making gcFront easy to use out-of-the-box, yet because it is a function in the MATLAB environment it can be easily integrated downstream of pathway designing tools, such as COBRA toolbox (Heirendt *et al*., 2019) and RetroPath2.0 (Delépine *et al*., 2018), and upstream of user-led design selection and robotics platforms, such as a CRISPR-Cas9-based KO platform done with an automated plasmid construction and transformation robot (Suckling *et al*., 2018) and automated adaptive laboratory evolution (Radek *et al*., 2017) – making gcFront a tool for the future of creating microbial cell factories.

## Supporting information

All Supplementary Information

## DATA AVAILABILITY

The data underlying this article are available in the article and its online supplementary information and data files.

## AUTHOR CONTRIBUTIONS

LL, AAM: developed theory; LL: developed code, ran procedures, plotted results; AAM, DGB: designed and supervised the research; all authors: wrote the paper.

