## Supplementary material for "gcFront: a tool for determining a Pareto front of growth-coupled cell factory designs": All Supplementary Information

### SUPPLEMENTARY FIGURES

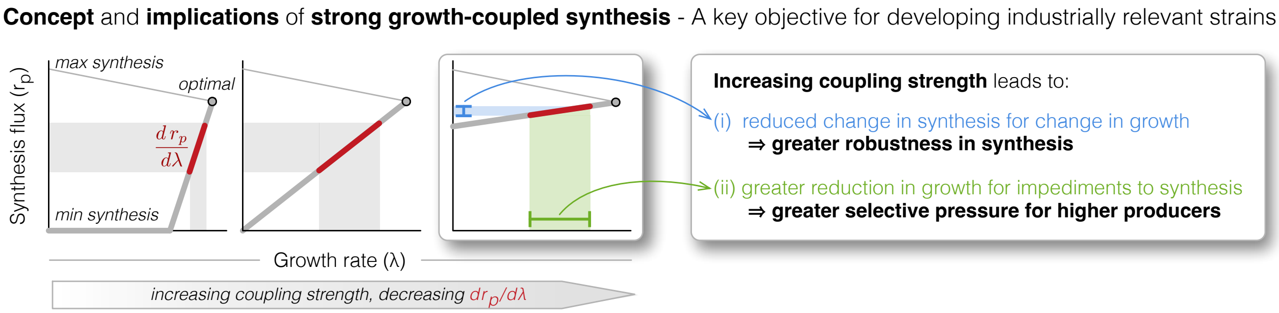

**Supplementary Fig. 1. Concept of growth-coupling and implications of selecting strong coupling strength.** Plots of example production envelops of strains with weak to strong coupling strength (from left to right). The plot of the production envelope is composed of the values of the minimum synthesis flux (thick lower grey line), maximum synthesis flux (thin upper grey line), and the optimal point (grey filled dot). For stronger coupling, the derivative of the change in synthesis for a change in growth (thick red line) is reduced. This has two implications: (i) synthesis is more robust to changes in drops in growth, e.g. suboptimal growths in large fermenters; and (ii) even for small impediments to synthesis (say via mutations over evolutionary time scales) there is a greater impairment to growth, which implies there is greater selection pressure for higher producers.

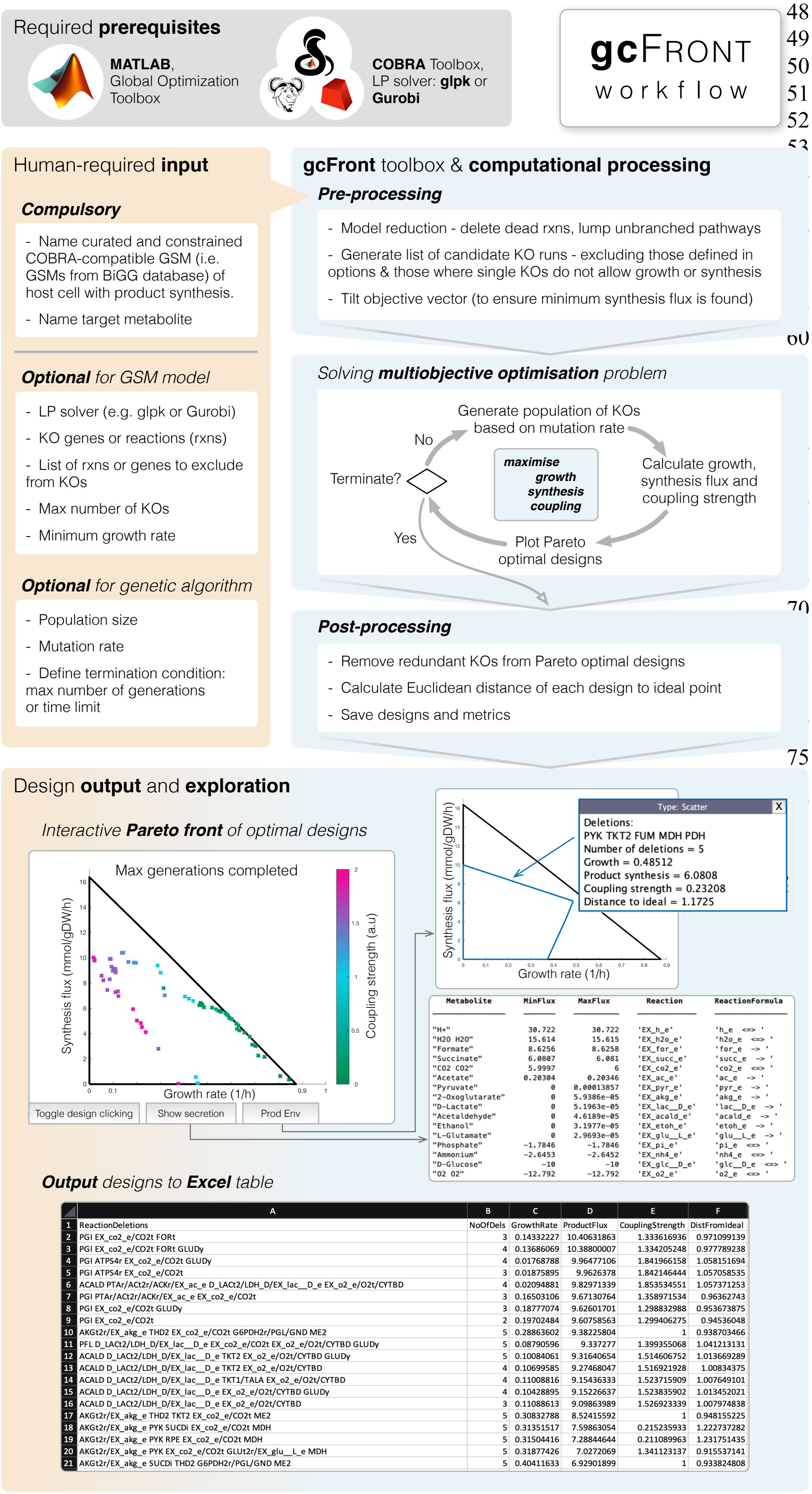

**Supplementary Fig. 2. The gcFront workflow.** A flow chart detailing (i) the required prerequisites for gcFront to run (gray box); (ii) the human required input of the COBRA-compatible and defined SBML model of the GSM of the cell host and product synthesis of interest, and options to adjust default parameters concerning GSM model and genetic algorithm (left orange box); (iii) what gcFront does "under-the-hood" (blue boxes), i.e. model reduction, iterative search for Pareto front of gc-designs that maximise growth, product synthesis flux, and coupling strength; and (iv) snapshots of the interactive plot of designs and performance metrics or option to save to Excel table (bottom box). See Supplementary Note 1 for details and gcFront documentation.

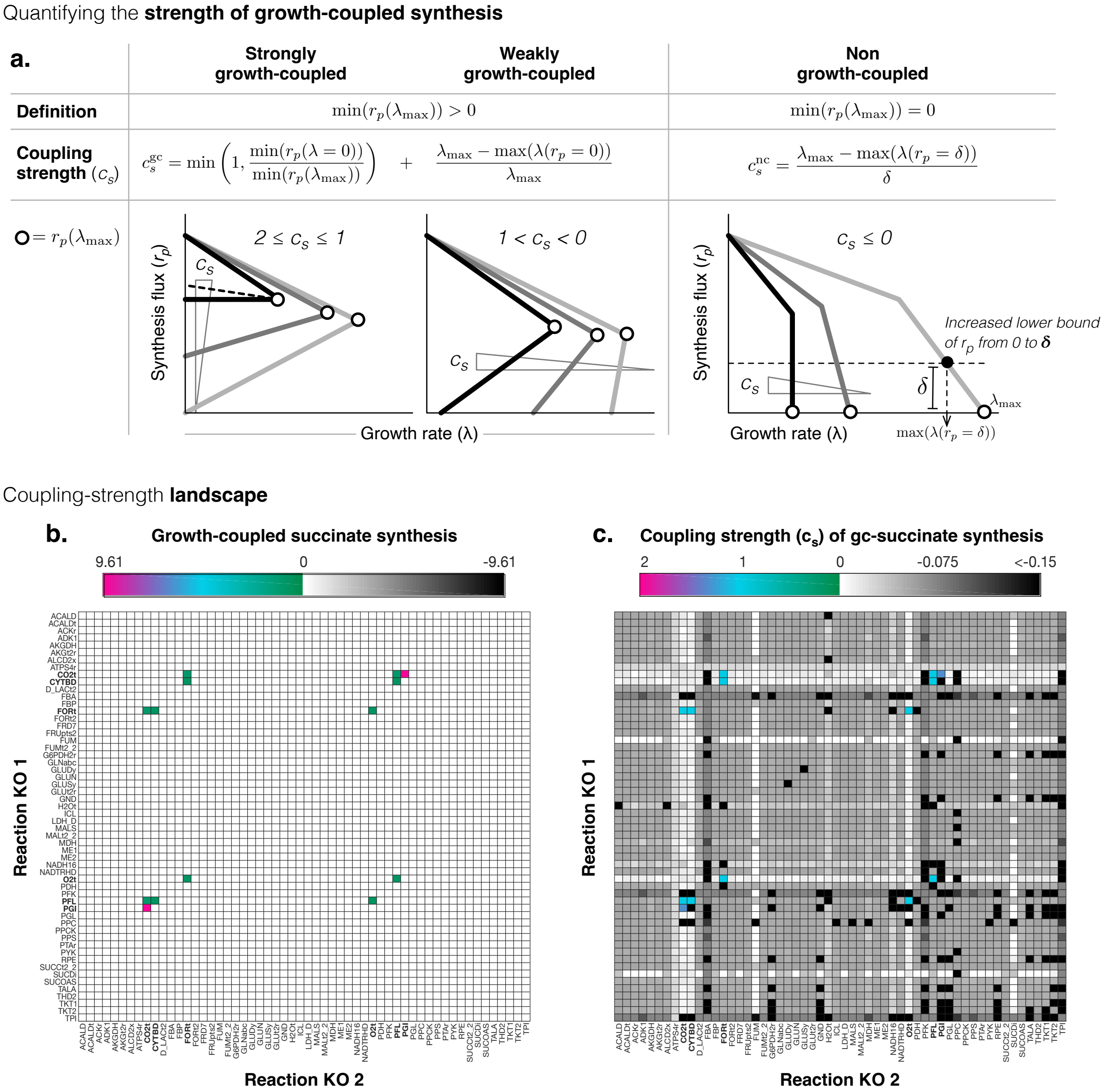

**Supplementary Fig. 3. Proposed measure of growth-coupling strength eases navigation of the design space.** (**a**). Illustration of the proposed coupling strength ($c_s$) measure, and how it is calculated from the production envelope, for designs (from right to left) that are non- , weakly-, and strongly-growth-coupled. (**b and c**). As an example, we explored pairs of reaction KOs to find growth-coupled (gc) succinate overproduction from gene-associated, non-essential reactions of the *E. coli* core GSM model (Orth *et al.*, 2010). For a search based on succinate synthesis, we observe a vastly flat landscape with rare spikes of gc-designs varying in succinate synthesis flux (**b**). Instead, scoring designs by $c_s$ could distinguish between non-gc designs (**c**, gray shaded boxes), creating a bumpier landscape that helps drive a bias to gc-designs (**c**, right plot, coloured boxes). We observe this bias in (**c**, right plot), where it tends to be the single KOs that yield non-gc designs with higher values of $c_s$ (lightest gray) that can be paired with another KO to yield a gc design (coloured boxes).

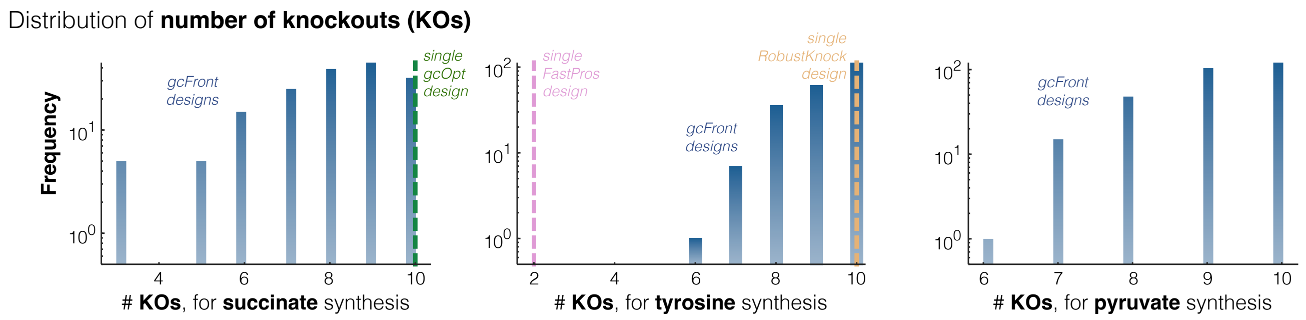

**Supplementary Fig. 4. Distribution of the number of KOs for growth-coupled designs**, of designs found (if any) from 6 hour runs of gcFront, the OptPipe implementation of RobustKnock, gcOpt, FastPros, and the COBRA implementation of OptGene (see Supplementary Notes 4 for details). gcFront and OptGene were run three times (N=3) to account for the stochastic nature of the genetic algorithm used for searching, so the concatenation of all designs found from those runs is shown here.

### SUPPLEMENTARY NOTES

#### Supplementary Note 1 - gcFront Documentation

##### *A. Preliminaries, dependencies, and installation*

We developed gcFront as a toolbox for use in MATLAB, but it is dependent on the following:

- MATLAB's Global Optimization toolbox for solving the multiobjective optimization problem.
- COBRA toolbox (Heirendt *et al.*, 2019) for the import and analysis of a COBRA-compatible GSM in SBML/XML format, such as those of the BiGGs database (King *et al.*, 2016). The full COBRA toolbox package can be installed from GitHub here: https://github.com/opencobra/cobratoolbox, and documentation and installation instructions can be found here: https://opencobra.github.io/cobratoolbox/stable/installation.html, and at their webpage here: https://opencobra.github.io/cobratoolbox/stable/.
- An LP solver for analysis of the GSM models, including flux balance analysis, flux variability analysis, and to enable gcFront to produce production envelopes, such as glpk or Gurobi (Gurobi Optimization, 2021) (recommended and free with an academic license) (downloadable from here https://www.gurobi.com/). This is a requirement of COBRA toolbox anyway, so any LP that successfully functions for the COBRA toolbox solver will also work for gcFront. The help guide for installation and testing of LP solvers with COBRA can be found in the section titled "Solver installation" in the COBRA installation webpage here: https://opencobra.github.io/cobratoolbox/stable/installation.html.
- gcFront was developed in MATLAB and with COBRA toolbox v3.1 and the LP problem was solved using Gurobi 8.1.1. gcFront was tested and successfully ran for different versions of MATLAB including MATLAB R2017a, R2018a, R2019a, R2020a and R2021a. We expect gcFront will function with other versions of these packages and LP solvers.

gcFront, with documentation and a workable tutorial, is freely available at GitHub: https://github.com/lLegon/gcFront, the repository of which is archived at Zenodo, DOI:10.5281/zenodo.6338595 (Legon *et al.*, 2022).

Once gcFront is downloaded, it is to be run using MATLAB. There are two approaches to run gcFront:

1. Change MATLAB's current directory to the gcFront folder downloaded and run it by typing "gcFront" into the command window.
2. Alternatively, add the gcFront folder to your MATLAB path, then you can call the function "gcFront" from within your own script or command window. The algorithm's inputs can be supplied by following the prompt windows that appear or can be supplied as arguments to the function.

##### *B. Overview of the gcFront algorithm*

See Supplementary Fig. 1 illustrating the gcFront workflow. gcFront is composed of four key steps: inputs, pre-processing of GSM, solving the multiobjective optimization problem, post-processing, and output of designs.

###### *(i) Inputs*

The gcFront function: [designTable, reducedModel, options] = gcFront(model, prodTarget, options).

Inputs:

- model = a genome-scale constraint-based model compatible with the MATLAB COBRA toolbox (XML/SBML). The model can be imported directly as a .mat file, or as a character array that specifies the file path to the model. If this input is left blank, users can browse to select the file from which the model should be loaded.
- prodTarget = Character array of the reaction or product/metabolite name in the GSM to be growth- coupled and overproduced. If a metabolite is specified, the target will be set to its exchange reaction.
- options = A MATLAB data structure with fields of the optional input parameter names, of the form "options.*Parametername*". The parameter names and default values are detailed in Supplementary Table 1, but

values of the parameters specified by the user will override the defaults. The order in which they are specified does not matter.

Outputs:

- designTable = a table of all Pareto optimal designs, including a column of the KO combination suggested, performance metrics (growth rate, minimum synthesis flux, coupling strength), and Euclidean distance to the ideal point normalized to maximum values to each objective value.
- reducedModel = MATLAB data structure of the COBRA GSM as reduced in the pre-processing.
- options = list of the values of the optional inputs used for this search.

**Supplementary Table 1. Optional inputs.** A list of all the optional inputs, with default values defined.

| Parameter name | Description | Default | Format |
|---|---|---|---|
| solver | The LP solver used to carry out FBA | Current COBRA toolbox LP solver. If none, the solver set by "initCobraToolbox" is used | Char, such as 'glpk' or 'gurobi', for example |
| biomassrxn | The name of the pseudoreaction that represents cell growth | Reaction with highest coefficient in objective vector | Char |
| tol | Tolerance to mathematical errors. Differences in flux smaller than this will be ignored | $10^{-8}$ | Positive double |
| tiltval | Coefficient used for objective vector tilting (so minimum product synthesis is identified) | $10^{-4}$ | Positive double |
| shiftval | Size of the change in growth rate that is used when manually calculating shadow price. | $10^{-5}$ | Positive double |
| skipreduction | Parameter that controls whether to reduce model size by removing inactive reactions and pooling linear pathways | False | Logical |
| mingrowth | Minimum growth threshold- designs with growth below this are ignored | $10^{-3}$ | Positive double |
| minprod | Minimum product threshold- any deletion that lowers product synthesis will not be considered when finding designs | $10^{-3}$ | Positive double |
| removeredundancy | Parameter that controls whether designs should be tested to see if they contain redundant deletions that do not contribute to the design | True | Logical |
| newredundantremoval | Parameter that controls whether redundant KOs are removed using the old method (as was used to generate data for this paper) or using a new, more efficient methodology. | True | Logical |
| maxreductionsize | Maximum number of KOs in designs that will have redundant KOs removed from them (helps avoid reduction of huge designs, which can take a long time). Not compatible with the old reduction method. | 15 | Positive double |
| saveresults | Defining if designs should be saved and exported to a csv file, and parameters are saved as .mat on termination | True | Logical |
| maxknockouts | Maximum number of knockouts that a design may have | Infinite | Positive integer double |
| deletegenes | Parameter that controls whether the algorithm searches for gene or reaction knockouts | False | Logical |

| | | | |
|---|---|---|---|
| ignorelistrxns | List of reactions that should be knocked out | Empty vector | Cell (contains characters) |
| ignorelistgenes | List of genes that should not be KO'd. If reaction KOs have been selected, reactions that can only be KO'd if these genes are KO'd are excluded from consideration | Empty vector | Cell (contains characters) |
| dontkoess | Parameter that controls if reaction KOs are ignored if they cannot be KO'd without a KO of a gene that is essential *in silico*. | True | Logical |
| onlykogeneassoc | Parameter that controls if reactions are only considered if they are gene associated | True | Logical |
| mutationrate | Average number of changes to a design during the GA mutation step | 1 | Positive double |
| popsize | Size of the GA population | 200 | Positive integer double |
| genlimit | Maximum number of generations that the GA will be run for | 10000 | Positive integer double |
| timelimit | Maximum number of seconds that the GA will be run for | 86400 (i.e. 1 day) | Positive double |
| fitnesslimit | GA will terminate if a design is found with a product synthesis that exceeds this value | Infinite | Double |
| spreadchangelimit | How low the change in the spread of designs must be before the algorithm starts to terminate. See MATLAB's 'gamultiobj' documentation for more details. | $10^{-4}$ | Positive double |
| stallgenlimit | How many generations the change in spread of the designs must be below spreadchangelimit before the algorithm is terminated | Same as genlimit (i.e. this will not lead to termination) | Positive integer |
| plotinterval | Number of generations that must pass before the plot of the current designs is updated | 1 | Positive integer |

***(ii) Pre-processing***

- First, all reactions in the model are subject to flux variability analysis (FVA) to see if they can carry non-zero flux, and are removed if they do not, i.e. are dead reactions.
- The algorithm then searches for reactions that form unbranched pathways and pools them into a composite reaction that represents flux through that pathway.
- If the algorithm has been set to search for gene knockouts, then the gene-reaction rules of the model are analysed to find sets of gene knockouts that are functionally equivalent, and to find sets of gene knockouts that can only have an effect if they are knocked out simultaneously. These genes are then pooled into their sets to reduce the size of the gene search space.
- Essential reactions are identified by determining single gene/reaction knockouts that reduce maximum growth rate or product synthesis flux below a user-defined minimum threshold ("mingrowth" parameter in Supplementary Table 1).
- A list of potential knockouts is then constructed, excluding essential, pooled, and dead reactions. Further reactions can also be excluded if the vector of cells listing those is added as the optional input parameter "ignorelistrxns", such as non-gene-associated reactions (including exchange reactions) or reactions that are disabled if an essential gene is knocked out, for example.
- Finally, the model objective vector (usually *modelname*.c in the COBRA model data structure) is "tilted", similar to (Feist *et al.*, 2010). Tilting is done by setting the value of the element of the c vector corresponding to the biomass reaction (growth rate) to +1, and the value of the element corresponding to the target reaction to a very

small negative value (-1*tiltval in Supplementary Table 1, e.g. $-10^{-4}$). This ensures that the product synthesis rate found by flux balance analysis that maximises growth rate is the minimum product synthesis rate.

***(iii) gcFront solving the multi-objective optimization problem***

Once executed, gcFront then moves to solve the multiobjective optimisation problem defined in Supplementary Note 2A, using MATLAB's Global Optimization toolbox function `gamultiobj`. The steps are as follows:

- An initial random population is created of size as specified in the optional input parameter "popsize" (Supplementary Table 1). Each member is defined by a different combination of KOs, as selected from the list of potential KOs generated during pre-processing, where KOs are implemented by fixing both the lower and upper bound flux constraints of those reactions to zero.
- Each member is then subject to flux balance analysis (FBA) and the three performance objectives are measured: growth rate, product synthesis flux, and coupling strength. Recall that the "tilted" objective vector determines both the maximum possible growth rate and minimum product synthesis flux possible at the maximum growth rate. The coupling strength ($c_s$) is calculated from the formula:

$$c_s = \begin{cases} c_s^{\text{nc}} = \dfrac{\lambda_{\max} - \max\left(\lambda(r_p = \delta)\right)}{\delta}, & \text{when } \min\left(r_p(\lambda_{max})\right) = 0, \\ c_s^{\text{gc}} = \dfrac{\lambda_{\max} - \max\left(\lambda(r_p = 0)\right)}{\lambda_{\max}} + \min\left(1, \dfrac{\min(r_p(\lambda = 0))}{\min(r_p(\lambda_{\max}))}\right), & \text{when } \min\left(r_p(\lambda_{\max})\right) > 0, \end{cases}$$

  as discussed in detail in Supplementary Notes 2B and illustrated in Supplementary Fig. 2a.
- Each population member is then ranked based on their dominance (i.e. if they have an equal or better score for every metric), and gcFront plots the designs of the full population of the current generation on the growth-synthesis plane, where each design is coloured by coupling strength. The production envelope of the parent model (i.e. with no deletions) is also plotted to help visualise how the growth and product synthesis of each design compares to that which is theoretically possible of the wild-type organism. If, however, no coupled designs have yet been discovered in the current generation, a scatter of growth rate vs coupling strength is plotted instead.
- A binary selection tourney is used to select designs of the population based on their rank and similarity to other designs, and those are then used to create new designs by mutation (adding/taking away random KOs) or crossover (taking random combinations of KOs from two parent designs).
- These steps are then repeated over many generations until the user specified termination condition is met (as defined in Supplementary Table 1), the default being whichever of "genlimit" or "timelimit" is hit first. Other optional termination conditions that can be set include "fitness limit", "spreadchangelimit", or "stallgenlimit".

***(iv) Post-processing and output of designs***

- Post-processing: On termination, the non-dominated gc-designs that make up the Pareto front are plotted and listed. Some of the optimal designs found may include KOs that are not contributing to performance. If desired (via optional input "removeredundancy"), gcFront removes such redundancy by testing each KO that makes up a design to see if the same growth, product synthesis and coupling strength can be achieved without it, removing the KO if it is not contributing to performance, and then repeating the process until no single deletion can be removed without moving the design off the Pareto front. As a simple metric to identify designs that lie in an optimal trade-off region, gcFront also measures the Euclidean distance ($D$) of the point of each design (defined by the three performance objective values) to the ideal point, after normalising it to the ideal point, i.e. $D = \sqrt{\sum_{i=1}^{3}\left(1 - \frac{\text{obj}_i^{\text{design}}}{\text{obj}_i^{\text{ideal}}}\right)^2}$. The ideal point lies at the point of maximum growth and maximum product synthesis rates, as calculated from FBA of the model with no deletions (the parent strain), and at the maximum possible coupling strength (i.e. 2).
- Output: All Pareto optimal gc-designs are then outputted in three formats simultaneously:
    - As a table onto the command window, listing all Pareto optimal designs, along with their KOs, the three performance measures, and distance to the ideal point.
    - As a table, exported to a .csv file, if the user specifies via optional input "saveresults".

- As an interactive plot on the growth-product synthesis flux plane, with variations in coupling strength illustrated as the variation in colors of designs (crosses), defined by the colorbar to the right of the plot. This plot helps visualise the inherent trade-offs between the objectives. Since this is an interactive plot, each design can be clicked to display their set of proposed KOs and performance metrics. Tabs at the bottom of the window also enable one to plot the production envelope of the respective design and a table of other side-products predicted to be secreted for that design.

For an example work-through, the users are referred to our tutorial in Supplementary Note 3.

#### Supplementary Note 2 - "Under the hood" of gcFront

##### *A. Identifying gc-designs - a multiobjective optimization problem*

The second key problem is to discover KOs that not only maximise coupling strength ($c_s$), as defined in Supplementary Note 2B, but also growth rate ($\lambda$) and product synthesis flux ($r_p$) - three fundamental performance objectives we look for in production strains. The growth-synthesis trade-off is understood (Varma *et al.*, 1993) and methods have been developed to elucidate this through multiobjective optimization (Zakrzewski *et al.*, 2012; Patané *et al.*, 2019; Andrade *et al.*, 2020), but consideration of the coupling strength is still lacking. We define the search for strain designs as a multiobjective optimization problem:

$$\max_{\underline{r}}(J_1, J_2, J_3),$$
$$J_1 = \lambda; \quad J_2 = r_p; \quad J_3 = c_s,$$
$$\text{subject to}$$
$$S \cdot \underline{r} = \underline{0},$$
$$\underline{k} \circ \underline{\text{lb}} \leq \underline{r} \leq \underline{k} \circ \underline{\text{ub}}, \text{ where } \sum_{i=1}^{R} k_i \geq R - I, \quad k_i \in \{0,1\},$$

for vector $\underline{r}$ of $R$ reaction fluxes of the GSM, excluding essential reactions, the biomass reaction, exchange reactions defining the growth media, and other user-specified reactions. $S$ is the stoichiometric matrix of the GSM (in reduced form after the preprocessing step), $I$ is the maximum allowable number of simultaneous knockouts (not an identity matrix), and $k$ is a vector where $k_i = 0$ defines KO of the $i^{\text{th}}$ reaction. This is implemented in the flux constraints as a Hadamard product of vector $\underline{k}$ and the vectors of the reaction fluxes' lower and upper bounds, $\underline{\text{lb}}$ and $\underline{\text{ub}}$. We solve this multiobjective optimization problem using the genetic algorithm function `gamultiobj`, from MATLAB's Global Optimization toolbox. Such metaheuristic methods have been shown to be less computationally demanding (Patil *et al.*, 2005; Rocha *et al.*, 2008), and although the solutions found cannot be guaranteed to be globally optimal this would have been computationally intractable anyway, given the massive search space.

It is important to note that if searching KOs over genes rather than reactions, the logical statement of how genes are associated to each reaction and a gene-reaction mapping matrix must exist in the GSM to enable KOs of the respective reactions to be made.

##### *B. A measure of coupling strength to ease the search for gc-designs*

To find KOs that predict growth-coupled synthesis from the GSM, we define these gc-designs as those that achieve a non-zero minimum product synthesis flux $r_p$ at the maximum possible growth rate $\lambda_{\text{max}}$, i.e. $\min\left(r_p(\lambda_{\text{max}})\right) > 0$, (Supplementary Fig. 2a). This is essentially a binary measure of gc, but creates a fundamental problem - KOs that enforce coupling appear as rare spikes in a vast flat landscape of non-gc-designs (Supplementary Fig. 2b, left), and the lack of driving bias towards a gc-design makes the search difficult and slow (Melo *et al.*, 2013). To distinguish between non-gc-designs, we define a measure based on the geometry of production envelopes generated from flux variability analysis i.e. phenotypic phase planes (Edwards *et al.*, 2002), similar to shadow price proposed in (Ohno *et al.*, 2014): $c_s^{\text{nc}} = \frac{\lambda_{\text{max}} - \max\left(\lambda(r_p = \delta)\right)}{\delta}$. This measure and its terms are illustrated in Supplementary Fig. 2a. In brief, to find $c_s^{nc}$: we perform two FBA calculations, the first to calculate the minimum synthesis flux at maximum growth rate (which will be zero for non-gc-designs) and the second to calculate the maximum growth rate after constraining the lower bound of the target reaction flux ($r_p$) to a small positive value $\delta$ (by default, $\delta = 1 \times 10^{-4}$), and calculate the ratio of the decrease in growth for the change in synthesis flux, as per the formula above. $c_s^{\mathrm{nc}} \leq 0$ for all non-gc-designs, but values closer to 0 indicate KOs that predict synthesis at a lower cost to growth (Supplementary Fig. 2a). This creates a bumpier landscape like that illustrated in Supplementary Fig. 2b (right). Interestingly, for that example plotted, we observe that single KOs where the product can be synthesised at lower growth costs (Supplementary Fig. 2c, right plot, lightest gray regions) can be paired with another KO to achieve coupling, demonstrating that $c_s^{\mathrm{nc}}$ can create a bias to find gc-designs more easily.

All gc-designs are not equal. Aiming to develop microbial factories where product synthesis is not only evolutionary favoured but robust to growth at suboptimal rates too, we define a measure of the strength of growth-coupled synthesis as $c_s^{\mathrm{gc}} = \frac{\lambda_{\max} - \max\left(\lambda(r_p = 0)\right)}{\lambda_{\max}} + \min\left(1, \frac{\min(r_p(\lambda=0))}{\min(r_p(\lambda_{\max}))}\right)$. This reduces the importance of weakly-coupled designs ($0 < c_s < 1$) where product is only guaranteed to be synthesized as maximum growth is approached, and favours strong coupling ($1 \leq c_s \leq 2$) where the product must be synthesised for the cell to grow at all, and is least affected for drops in growth rate. Altogether, our definition of coupling strength creates a continuous measure

$$c_s = \begin{cases} c_s^{\mathrm{nc}}, & \text{when } \min\left(r_p(\lambda_{\max})\right) = 0, \\ c_s^{\mathrm{gc}}, & \text{when } \min\left(r_p(\lambda_{\max})\right) > 0, \end{cases}$$

to help find KO designs that are strongly coupled.

#### Supplementary Note 3 - Tutorial for finding gc-designs

##### *A. Work-through example - finding gc-design for succinate overproduction in E. coli core*

1. First, ensure that all prerequisites (MATLAB Global Optimization toolbox, COBRA toolbox, LP solver) are available or have been downloaded and are working correctly (gcFront should warn you if they are not).

2. Also, ensure that you have added the gcFront_code folder to your MATLAB path. This only needs to be done the first time. To do this go to the home tab and select “Set Path” under the environment panel, like so:

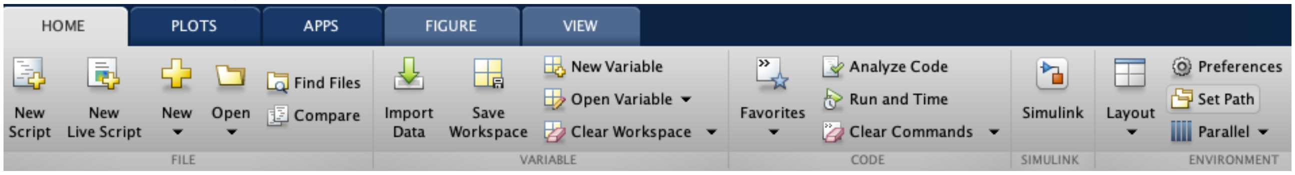

Then click “Add Folder…”, navigate to the gcFront folder, click “Open”, and finally click “Save”:

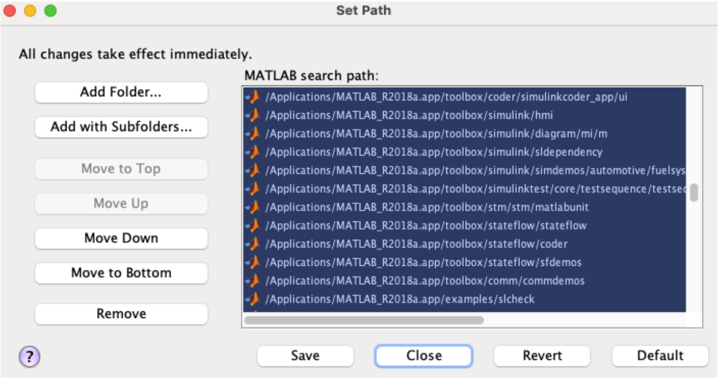

3. Then, download the model of interest. In this example, we will use the *E. coli* core GSM model (Orth *et al.*, 2010) from the BiGG model database (King *et al.*, 2016):

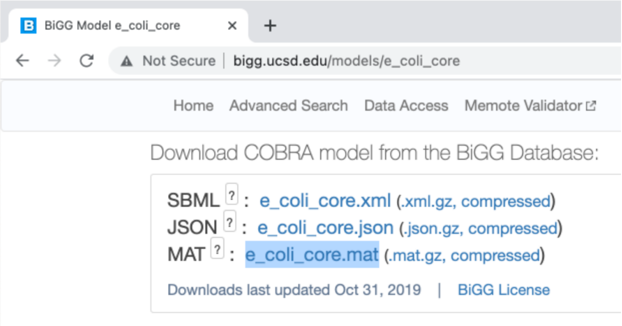

4. Before running gcFront, we should ensure all model reaction constraints are set for the defined media condition, genotype, etc, and save the changes. For this *E. coli* core model, we use the COBRA toolbox to set the maximum glucose uptake rate to 10 mmol/gDW/h by adjusting the lower bound of the glucose exchange reaction to -10 mmol/gDW/h:

```
% setting glucose supply to 10 mmol/gDW/h in E. coli core model
% by changing lower bound of exchange reaction
model=readCbModel('e_coli_core.mat');
model=changeRxnBounds(model, "EX_glc__D_e", -10, 'l');
save('e_coli_core.mat','model');
```

5. Type "gcFront" into the command window (if desired, algorithm inputs and parameters can be supplied as arguments to the function as well - see the SUCCINATE_TUTORIAL.m file in the code downloadable from GitHub for an example of how to do this).

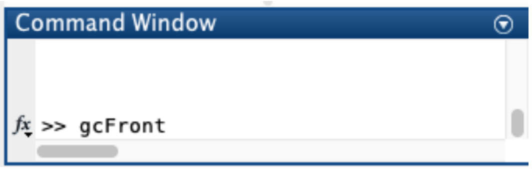

6. Enter the file path and name of the GSM that needs to be imported (here it is "e_coli_core.mat"), and the reaction or metabolite that you want to couple (here it appears as "succinate" in the GSM):

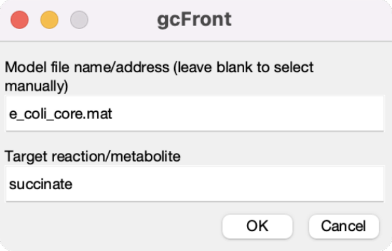

7. After clicking OK, another window appears. Here we can redefine optional input parameters with the drop-down menu, defining the new values in the lower text box. Each parameter is assigned a default value, but the default is overridden by any user-defined value. Here, we set the maximum number of KOs to 5 and enable results of all Pareto optimal designs and their performance measures to be outputted into a table in a csv file by setting "saveresults" to true or 1:

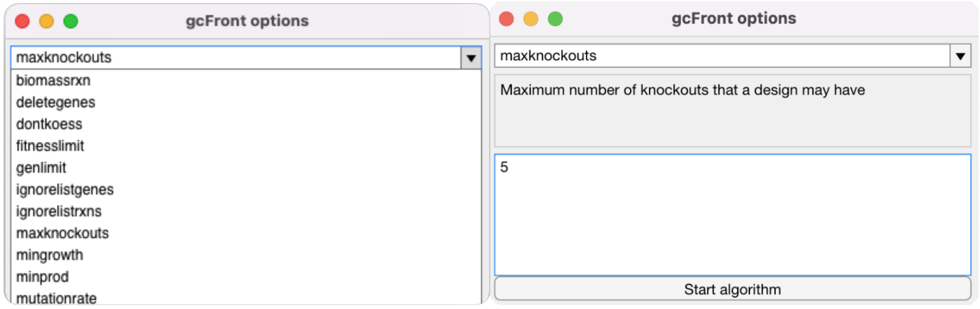

8. After clicking on “Start algorithm”, gcFront will then begin - it iteratively solves the multi-objective optimization problem, updating a plot of all the designs of the current generation on a 2D plot of the objective values (i.e. target flux vs growth rate, with designs (dots) coloured by coupling strength).
At first, it may plot the growth rates and coupling strengths for the non-gc-designs identified, as shown here:

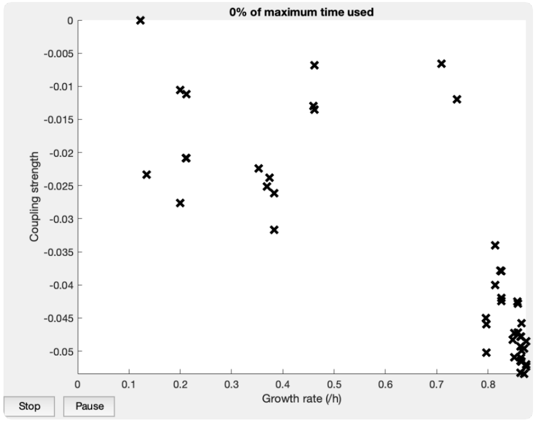

As soon as gc-designs are found, the plot is updated to display growth rates (x-axis), product synthesis flux (y-axis), and coupling strengths (colour of each point), of all designs of the current generation, together with the production envelope of the parent genotype (model with no deletions). Users can end the optimization early by pressing the “Stop” button as shown here:

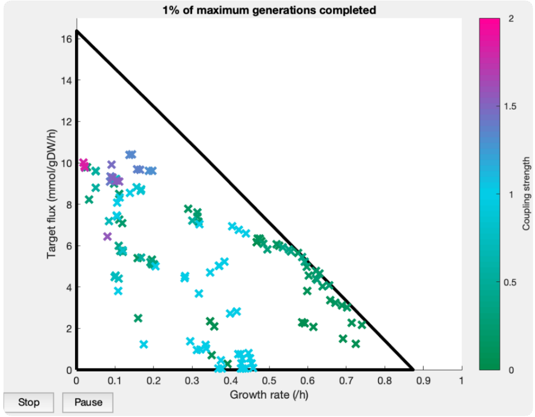

9. Once the algorithm is stopped or reaches a termination condition, only the Pareto front of the final population of designs are displayed on the plot and in the command window. Designs will also be saved as a .csv file in the current folder since the “saveresults” option is enabled by default.

A slash in the name of a deletion indicates that any one of the genes/reactions can be KO’d to achieve the specified performance (so in the example below, the performance of the design on the first line could be achieved with a KO of the reactions PFL, PGI and either EX_co2_e or CO2t).

| ReactionDeletions | NoOfDels | GrowthRate | ProductFlux | CouplingStrength | DistFromIdeal |
|---|---|---|---|---|---|
| "PFL PGI EX_co2_e/CO2t" | 3 | 0.14332 | 10.406 | 1.3336 | 0.9711 |
| "PFL PGI EX_co2_e/CO2t GLUDy" | 4 | 0.13686 | 10.388 | 1.3342 | 0.97779 |
| "PFL D_LACt2/LDH_D/EX_lac__D_e ETOHt2r/ALCD2x/EX_etoh_e THD2 EX_o2_e/O2t/CYTBD" | 5 | 0.017631 | 10.025 | 1.8369 | 1.057 |
| "ACALD D_LACt2/LDH_D/EX_lac__D_e THD2 EX_o2_e/O2t/CYTBD" | 4 | 0.090648 | 9.9108 | 1.4837 | 1.0129 |
| "PFL D_LACt2/LDH_D/EX_lac__D_e ETOHt2r/ALCD2x/EX_etoh_e EX_o2_e/O2t/CYTBD" | 4 | 0.020154 | 9.8848 | 1.8488 | 1.0571 |

An interactive plot of the Pareto front of the discovered designs will also be displayed. Users can click on designs to display information about them:

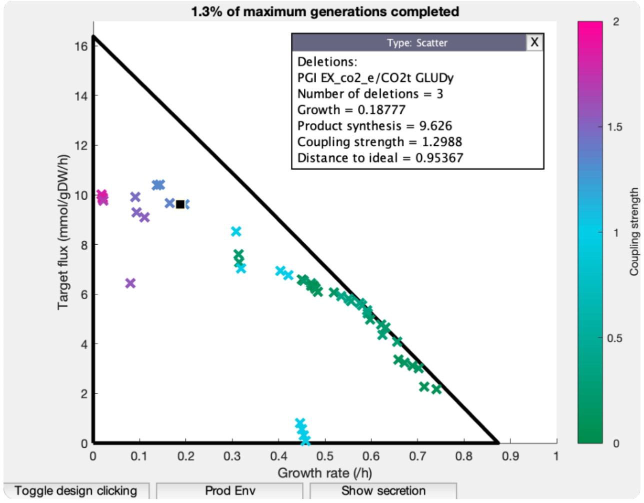

Also, users can use the "Prod Env" button to display the production envelope of the selected design, and use the "Show secretion" button to show predicted metabolite secretion/uptake for the selected design:

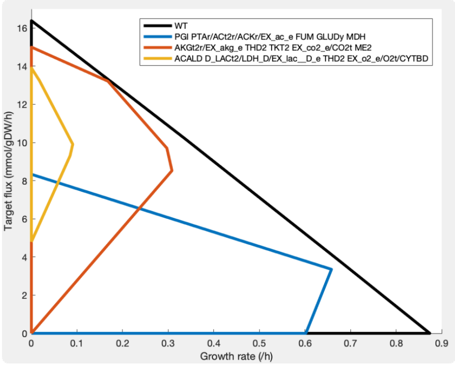

Deletions: PGI EX_co2_e/CO2t GLUDy

| Metabolite | MinFlux | MaxFlux | Reaction | ReactionFormula |
|---|---|---|---|---|
| "H+" | 32.324 | 32.324 | 'EX_h_e' | 'h_e <=> ' |
| "H2O H2O" | 10.994 | 10.994 | 'EX_h2o_e' | 'h2o_e <=> ' |
| "Succinate" | 9.626 | 9.6261 | 'EX_succ_e' | 'succ_e -> ' |
| "Formate" | 5.1059 | 5.106 | 'EX_for_e' | 'for_e -> ' |
| "Acetate" | 4.1995 | 4.1997 | 'EX_ac_e' | 'ac_e -> ' |
| "Pyruvate" | 0 | 0.00010937 | 'EX_pyr_e' | 'pyr_e -> ' |
| "D-Lactate" | 0 | 4.1015e-05 | 'EX_lac__D_e' | 'lac__D_e -> ' |
| "Acetaldehyde" | 0 | 3.6458e-05 | 'EX_acald_e' | 'acald_e -> ' |
| "2-Oxoglutarate" | 0 | 2.9829e-05 | 'EX_akg_e' | 'akg_e -> ' |
| "Ethanol" | 0 | 2.524e-05 | 'EX_etoh_e' | 'etoh_e -> ' |
| "L-Glutamate" | 0 | 1.6406e-05 | 'EX_glu__L_e' | 'glu__L_e -> ' |
| "Phosphate" | -0.69075 | -0.69075 | 'EX_pi_e' | 'pi_e <=> ' |
| "Ammonium" | -1.0239 | -1.0239 | 'EX_nh4_e' | 'nh4_e <=> ' |
| "O2 O2" | -7.1489 | -7.1489 | 'EX_o2_e' | 'o2_e <=> ' |
| "D-Glucose" | -10 | -10 | 'EX_glc__D_e' | 'glc__D_e <=> ' |

10. Users must then choose between these designs to find ones that are best suited to their requirements. Alternatively, they can develop bespoke criteria or measures to determine designs that balance between the objectives as they deem suitable. More details of this in the next subsection *(B)*.

##### *B. Examples of ways to harness the results to select gc-designs to take to the lab*

gcFront produces a number of designs. To select between these, users must make decisions about the trade-offs between growth, product synthesis, and coupling strength that they are willing to accept. A high growth rate results in faster synthesis processes and raises productivity, high synthesis rates raise both yield and productivity, and a stronger coupling strength can encourage stronger selective pressure for product synthesis and robustness of synthesis to suboptimal growth rates. After choosing a design with favourable parameters, users should assess the biological feasibility of this design if this was not already done at an earlier stage. If toxic metabolite production was not constrained in the model, then users should check what byproducts their coupled designs are expected to result by clicking on the "Show secretion" button. If substantial toxic byproduct secretion is predicted, the user should then check whether their design is still functional after constraining the secretion of the toxic byproduct. In addition, if gcFront was not used to search for gene knockouts, then users must identify genes affiliated with their reactions to convert their reaction knockouts into a strategy that can be experimentally implemented. Furthermore, if essential gene knockouts were not already excluded from consideration, then biological knowledge should be used to assess the feasibility of the identified gene knockouts, as some genes that are not essential *in silico* cannot be knocked out experimentally.

#### Supplementary Note 4 - Running other gc-designing algorithms

***How each algorithm was run***

We tested the time to find the first gc design and the gc designs found by each algorithm, without the need for high process computing. To ensure a fair comparison, each algorithm was run in MATLAB 2017a on a laptop (MacBook Pro with 2.3 GHz Quad-Core Intel core i5 processor, 8GB 2133 MHz LPDDR3 RAM), with Gurobi (Gurobi Optimization, 2021) used as the LP solver (where possible), until a design was discovered, or for a maximum of 6 hours if no design was discovered in this time. If an algorithm terminated before 6 hours had passed because a gc-design was found, or due to premature termination, it was then run for a full 6 hours and the designs found by the end were saved, if any. Each algorithm was also run using the same *E. coli* GSM iML1515 model with its default constraints to determine reaction KOs for gc-synthesis of succinate, L-tyrosine, and pyruvate. To try and limit solutions to more biologically feasible designs, reaction KOs were only considered if the reaction was gene-associated and non-essential, based on analysis of the GSM model or found experimentally (Goodall *et al.*, 2018). We ran each algorithm essentially "out-of-the-box" as best we could, but some minor modifications had to be made to allow for them to terminate after the discovery of a gc-design, and to ensure they only looked for knockouts of the non-essential reactions. These modifications should not affect the designs identified, but we discuss the modifications made for running each procedure below.

***A. RobustKnock***

The original implementation of RobustKnock (Tepper and Shlomi, 2009) depends on TOMLAB (which does not currently offer free academic licenses), so we performed RobustKnock based on its implementation in OptPipe (Hartmann *et al.*, 2017). The "irrelevantRxns" function was modified so essential reactions could be excluded from further consideration, and the "testRxnDeletion" function was modified so no further reaction deletions would be tested either after the 6 h time limit had passed, or once a growth-coupled design had been discovered. This ensured swift termination after the discovery of a gc-design.

***B. gcOpt***

gcOpt (Alter and Ebert, 2019) uses Gurobi to solve a MILP problem. The gcOpt code was modified so it would supply Gurobi with the "BestObjStop" parameter, which terminates the optimisation process after the objective exceeds a given threshold. In this way, it became possible to terminate the algorithm once a gc-design was discovered, and so determine the minimum time required to discover a gc-design.

***C. FastPros***

FastPros (Ohno *et al.*, 2014) normally considers gene-reaction associations when making knockouts, so the model was modified so that each reaction of interest was listed as being associated with a single unique gene, while all other reactions were listed as not being gene associated. By doing so, FastPros was forced to consider reaction knockouts, making for a fairer comparison. To allow for termination of the algorithm, the "calcUtargetSelKoStrains" function was modified so that no further knockouts were considered after the time limit was exceeded, and postprocessing was only conducted on knockouts with a shadow price that indicated that they might be growth coupled. In addition, the "FastPros" function was modified so the algorithm terminated after discovery of a gc-design.

***D. OptGene***

The original OptGene was implemented in C++ (Patil *et al.*, 2005), so we chose to use the MATLAB implementation of OptGene from the COBRA toolbox (Heirendt *et al.*, 2019) for a fairer comparison. To enable the algorithm to terminate once a design with a non-zero product synthesis flux is discovered, the "optGene" function was modified so that the options that it passes to MATLAB's genetic algorithm function (ga) included a "FitnessLimit" parameter, and the "GetOptGeneSol" function was modified so it does not return an error if a gc-design was not discovered within the 6h time limit.
